# Transformation of Quotient Values for their use as Continuous Cladistic Characters

**DOI:** 10.1101/333781

**Authors:** Steven Uros Vidovic

## Abstract

Recent advances in cladistic technology have produced novel methods for introducing morphological data into cladistic analyses, such as the landmark and continuous character functions in the software TNT and RevBayes. While these new methods begin to address the problem of representing morphology, there has been little consideration of how to transform and code the operational taxonomic units’ (OTUs) dimensions into the datamatrix. Indeed, angles, serial counts, percentages and quotient values can be used as continuous characters, but little has been said about how coding these data affect the trees discovered. Logically, counts of elements and angles measured off specimens may be coded directly into continuous character matrices but percentages and quotient values are more problematic, being transformed data. Quotient values and percentages are the simplest way of representing proportional differences between two dimensions and reducing the effect of inter-taxonomic magnitude differences. However, both are demonstrated to be problematic transformations that produce continuous characters with weighted states that are non-representative of morphological variation. Thus, two OTUs may be represented as less/more similar morphologically than other OTUs that display the same degree of morphological variation. Furthermore, the researcher’s choice of which dimension is the divisor and dividend will have a similar affect. To address this problem, a trigonometric solution and a logarithmic solution have been proposed. Another solution called linear transposition scaling (LTS) was recently presented, with the intention of best representing and coding observable morphological variation. All three methods are reviewed to establish the best way to represent and code morphology in a cladistic analysis using continuous characters.

---

What is the best way to represent and code morphology for cladistic analysis? “The holy grail” (Thiele 1993) of morphological cladistics is the inclusion of continuous characters for the representation of morphometric data. Consequently, many methods of continuous (quantitative) data inclusion have been proposed, including gap coding (Mickevich and Johnson 1976); gap weighting (Thiele 1993); finite mixture coding (Strait et al. 1996); and overlap coding (Swiderski et al. 1998). Recent advances in representing morphological data for cladistic analysis have been the inclusion of landmark (Goloboff and Catalano 2016) and continuous character functions in the software TNT (Goloboff et al. 2008b). Despite concerted efforts to represent morphology in cladistic analyses using continuous data, there has been little discussion about coding, *a priori* data transformation, or the possible weighting effects of including this type of data (Wiens 2001).

However, these data cannot be ignored. Ignoring morphological data in phylogenetic studies means ignoring the fossil record (Hillis 1987). Fossil data can both resolve phylogenetic relationships for clades represented only by extinct taxa, and inform and complement molecular studies, helping to resolve internal node relationships for crown groups. Additionally, the accurate phylogenetic placement of fossil taxa is the best way to time-calibrate trees for chronological (clock and rate) studies.

The size and shape of organisms is fundamental to the analysis of biological variation (Jolicoeur and Mosimann 1960). Methods for expressing similarity of shape and relative dimensions quantitatively have been extensively reviewed over the last half a century (Jolicoeur and Mosimann 1960; Mosimann 1970; Kendall 1981; Bookstein 1982; Kendall 1984; Bookstein 1989; Small 1996; Rohlf 1998, 2003; Adams et al. 2004; Baur and Leuenberger 2011). “Many, if not most, morphological characters describe variation in quantitative traits (e.g. differences in size, shape, or counts of serially homologous structures), regardless of whether systematists choose to code them quantitatively orqualitatively (Stevens 1991; Thiele 1993)” (Wiens 2001). Certainly, ratios, quotient values, percentages, serial counts, angles and other dimensions are commonly used in taxonomic diagnoses and cladistic analyses, often providing the only significant separation of cryptic taxa (e.g. Vidovic and Martill 2014) that mostly lack distinct qualitative characters (Baur and Leuenberger 2011). As such, measurements of dimensions *a* and *b* expressed as ratios (a:b) or quotient values (a/b) are entrenched in systematic literature. This study is primarily concerned with these simply transformed quotient values as an expression of two-dimensional morphological variation, and their use as continuous cladistic characters.

The effects of representing morphological variation as quotient values is by no means limited to maximum parsimony analysis. Preliminary experimentation demonstrates the effects of the transformations discussed here on out to result in the recovery of incongruent tree topologies from Bayesian inference in RevBayes too. Because RevBayes and TNT search for trees using the data matrices in vastly different ways, the focus of this study is on maximum parsimony and specifically analyses conducted in TNT.

The continuous character function in the maximum parsimony software, TNT, works by applying a weight between 0.000 and 65.000 to values within that range. Thus, it is possible to have sixty-five thousand individual states and the more similar two taxa are, the fewer steps implied on a tree. Whereas the less similar two OTUs are, the greater number of steps implied. Several problems result from this continuous method: 1) the sum of parts (Pereyra and Mound 2009) can cause a single continuous character to have a far greater influence over a tree search than any discrete equal weights characters; for quotient values specifically: 2) ‘converse data’ (i.e. where one dimension is greater than another in one OTU, but the opposite is true in another OTU) (Fig. 1) will cause a bias towards lumping OTUs that have a smaller dividend than the divisor and splitting of OTUs demonstrating the converse (Vidovic and Martill 2014); 3) depending on which dimension a researcher choses to be their dividend or divisor, they will get different results (Mongiardino Koch et al. 2015).

**Figure 1.**
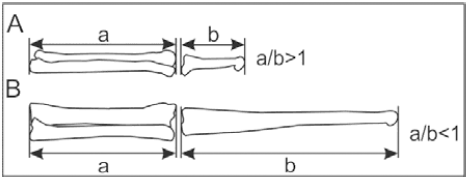
Two reconstructions of pterosaur radii, ulnae (a) and wing metacarpi (b), demonstrating a converse relationship between the dividend and divisor of taxa closest to *Eudimorphodon* (A) and taxa closest *Pteranodon* (B).

It is possible to transform data *a priori*, to counteract these problematic features of continuous characters. (1) Pereyra and Mound (2009) suggested normalising data to a range that would imply more suitable weights for the states. A (2) trigonometric transformation (Vidovic and Martill 2014) and a (3) logarithmic transformation (Mongiardino Koch et al. 2015) of quotient values prior to their inclusion in cladistic analyses have also been suggested. Vidovic and Martill (2014) used a trigonometric solution to transform data that demonstrated a converse relationship among OTUs in a cladistic analysis. In practice, this meant that where magnitude of element *a* exceeded element *b* in one OTU, and element *b* exceeded element *a* in another OTU, the equation *i*=*tan*^*-1*^*(a/b)* was applied to the data. An example of morphological attributes affected by converse data is the length of the pterosaur wing metacarpal (MCIV) relative to the ulna (Fig. 1), where MCIV can be shorter than the ulna in species most like *Eudimorphodon* (Fig. 1A), or longer in species most like *Pteranodon* (Fig. 1B). These converse relationships can be seen throughout vertebrate and invertebrate anatomical variation in homologous structures. Other examples include, the length/depth of bird bills, the length/width of spider abdomens, and in the length/width of elements in the appendicular skeletons of secondarily marine reptiles and fossorial vertebrates. These occurrences are by no means rare in nature.

Mongiardino Koch et al. (2015) noted a different, but similar scenario to Vidovic and Martill (2014) that affected the results of cladistic analyses and proposed a logarithmic transformation of quotient values. According to Mongiardino Koch et al. (2015), depending on which value is chosen by the researcher to be the dividend or divisor the outcome of the cladistic analysis will be different. The reason for the same data producing different results is that calculating a quotient value greater than unity does not transform the input data in the same way as calculating a quotient value less than unity (Fig. 2A & B; Fig. 3). The effect identified by Mongiardino Koch et al. (2015) is the same as the one demonstrated by Vidovic and Martill (2014: Figure 5), but researcher generated.

**Figure 2.**
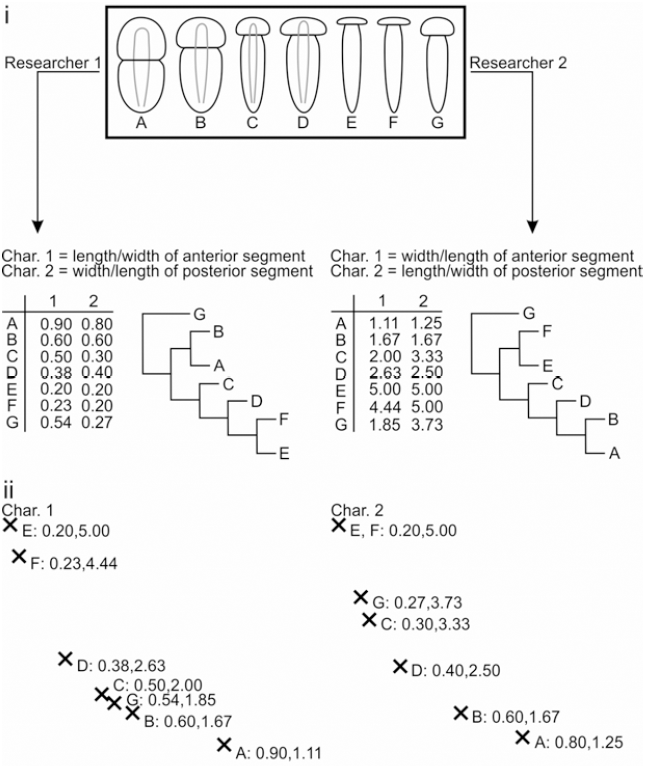
i) Adapted from Mongiardino-Koch et al. (2015) – expanded upon. Hypothetical organisms A–G are studied by two researchers, Researcher 1 and Researcher 2. Both researchers study the same dimensions, but calculating their quotient values as the inverse of the other’s. Researcher 1 finds a different most parsimonious tree to Researcher 2, despite seemingly treating the data the same. ii) Character 1 states used by Researcher 1 (x-axis) are plotted against those used by Researcher 2 (y-axis) and character 2 is plotted in the same way. These plots demonstrate that by dividing a smaller number by a larger number, the states analysed by Researcher 1 are on an exponential scale.

**Figure 3.**
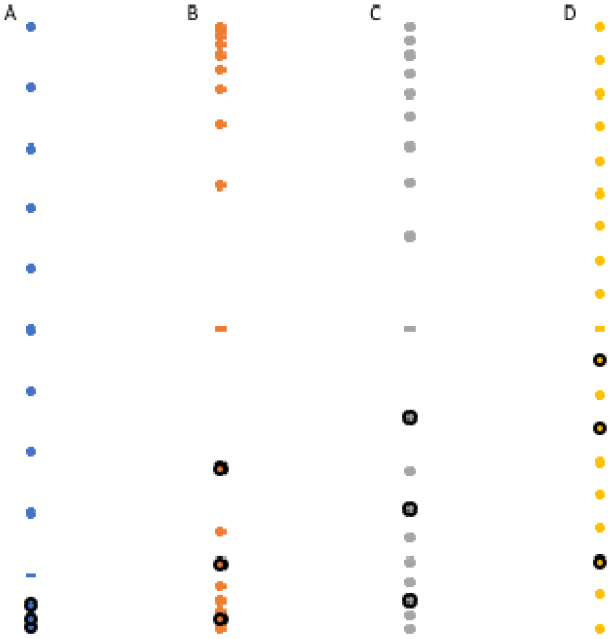
A) Graphical representation of contiguous integers 1–10 and their inverse forms(i.e. 1 divided by 2–10) and those values transformed using the: B) trigonometric method; C) logarithmic method; and D) the LTS method. The dashes represent unity, which is the centre of the total range of disparity, and the ringed points are the functions of 2, 4 and 8. Note that the proximity of these points on the graph represent the relative weight difference and therefore similarity/dissimilarity as perceived by a cladistic analysis.

Logarithmic and trigonometric transformations result in weights being applied to OTUs that determine the same weight difference between two OTUs if they demonstrate one morphological variation or the directly inverse relationship. However, the two methods do not produce the same states (weights) with the same source material and if the data is normalised to the same range. Thus, the tree topology recovered by a cladistic analysis will be affected by the transformation that is chosen. Furthermore, another method termed linear transposition scaling (LTS) was presented by Vidovic (2016) and Vidovic and Martill (2017). LTS also produces different results to logarithmic and trigonometric transformations, and presently it is not clear which transformation best represents and codes continuous data. Here, the effect of using different quotient value transformations (logarithmic, trigonometric and LTS) on the data distribution and therefore cladistic continuous character weights is discussed with examples. The example of Mongiardino Koch et al. (2015) using Anna and John is discussed further to support the concept of LTS. Additionally, graphical and cladistic experimentation is used to compare the methods in question and test the validity of using quotient values to represent a simple morphometric condition by comparing the LTS method to geometric morphometric principal component analyses.

## Example

Mongiardino Koch et al. (2015) proposed a hypothetical situation where two researchers, Anna and John, made different decisions when writing characters for morphometric data available from their studied taxa. Both Anna and John were testing ingroup relationships for the genus A+B+C+D, with the outgroup E. Here, in an adapted example, Researcher 1 and Researcher 2 take the OTUs studied by Anna and John (A, B, C, D, and E) and include additional OTUs F and G. Researchers 1 and 2 both proposed continuous morphometric characters that would study the dimensions of the anterior and posterior segments of their taxa. The results of their decision-making processes are, Researcher 1 had quotient values less than one, whereas Researcher 2 had quotient values greater than one (see the tables in Fig. 2ii). One may expect both analyses to recover the same tree, given that they are analysing the same morphometric data and treating it seemingly in the same way. However, Researcher 2’s result proposes a different phylogenetic hypothesis to that of Researcher 1. The reason for the alternative phylogenetic hypotheses is that it costs less for each tree respectively, due to the data transformations distributing the states differently and implying different weights. Note, states for characters 1 and 2 plotted as Researcher 1 vs Researcher 2 (Fig. 3ii) do not have a directly linear relationship.

Mongiardino Koch et al. (2015) assumed that neither Anna nor John were right or wrong to treat the data the way they did. However, either Anna or John are proposing phylogenetic hypotheses contradicting the exact phylogeny more than the other.

On closer inspection of the characters used by Researcher 1 and Researcher 2 it is clear that Researcher 1’s morphological ranges become exponentially larger as their quotient values approach one (Fig. 2ii). Mongiardino Koch et al. (2015) tested several data transformations, including range scaling and z-scoring, attempting to eliminate the effect of researcher decision. However, they did not reason whether or not Anna or John had provided the correct result. In effect, they wanted any dividend/divisor relationship to produce the same result, assuming that would be the correct result. To resolve scale inequality Mongiardino Koch et al. (2015) proposed that the data should be log transformed. Similarly, Vidovic and Martill (2014) proposed a trigonometric transformation method. Although both methods solve scale inequality, they too are on exponential scales and have an undesirable impact on the entire dataset.

In conclusion, Researcher 2 is representing and coding the data with greater fidelity than Researcher 1, due to Researcher 2 unwittingly making the research decision that best represented the source data as weighted states (Fig. 2ii). Therefore, when calculating quotient values, it is advisable to have the larger number as the dividend and the smaller number as the divisor. Keeping that in mind the LTS method has been developed and is described and tested below.

## Materials and Methods

### Rationale

Two qualities are required of the states resulting from the transformation of morphometric data for them to be effective continuous character states. These criteria are defined as follows:

1. Multiplying the morphological variation by N must increase the weight implied (i.e. the state) by the same multiplier (N). Likewise, adding a unit to a dimension N times must increase the weight implied the same amount N times.
2. Values need to be interchangeable for the dividend and divisor, producing the same distribution of states either way.

These criteria are simply tested by observing univariate plots of quotients and transformed quotients where ten of the calculations *a* is a value between one and ten (serially increasing by a single unit), and *b* is one. The remaining nine calculations are inverted i.e. *a* is one and *b* is a value of two to ten. The values 2, 4 and 8 are chosen to observe if the first tenet of criteria 1 is true of each transformation/treatment of the data. The general distribution of the plots is used to infer the validity of the second tenet of criteria 1, although the tenets are co-dependent. On the plots, unity is marked with a dash to help interpretation of criteria 2.

### Real Data Examples

To test the reliability of continuous data transformed using a logarithmic transformation, trigonometric transformation and LTS, two experiments were conducted. The Pterodactyloidea was used to test the morphometric cladistic techniques because it is a morphologically well-defined and constrained group; the group is represented in the fossil record from their appearance to extinction; only morphological characters are available for this group; they demonstrate inverse proportions between taxa (Fig. 1). The analysis of Vidovic & Martill (2014) was chosen because compound characters – which have been demonstrated to be detrimental to optimal tree recovery (Brazeau 2011) – have been reduced; it is the most comprehensive pterodactyloid cladistic analysis currently in the literature; currently it is the only pterodactyloid cladistic analysis to utilize continuous characters.

#### Method one

Data from one continuous cladistic character (Vidovic and Martill 2014: Ch. 1) – demonstrating a converse relationship between the dividend and divisor in the range of taxa being studied – is plotted on a graph (Fig. 4). The distribution of untransformed (Fig. 4A) and transformed (Fig. 4B-D) quotient values were compared graphically (univariate plot) for a/b; b/a; and a/b inverted so that direct comparisons of the data distribution could be made visually. All results, including previously untransformed data, were normalised between naught and one for comparison with each other and to avoid the character net weight issue identified by Pereyra and Mound (2009). In this case the character being analysed is the rostral index, using the data of Vidovic and Martill (2014: Ch. 1). Therefore, the distributions of rostrum depth/length and rostrum length/depth for pterodactyloid pterosaurs were compared for datasets using no data transformation (Fig. 5A), and LTS (Fig. 4B), trigonometric (Fig. 4C) and log ratio (Fig. 4D) transformations.

**Figure 4.**
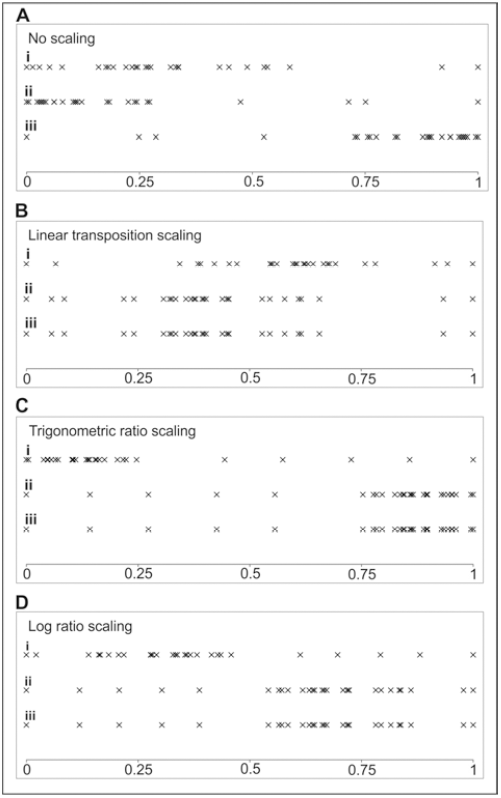
Univariate plots of character states taken from Vidovic and Martill (2014) character 1 and transformed using different methods: A) no transformation; B) LTS; C) trigonometric transformation; and D) logarithmic transformation. For each method, the character states for i) rostrum depth/length, ii) rostrum length/depth, and iii) depth/length in reversed order are plotted for comparison.

**Figure 5.**
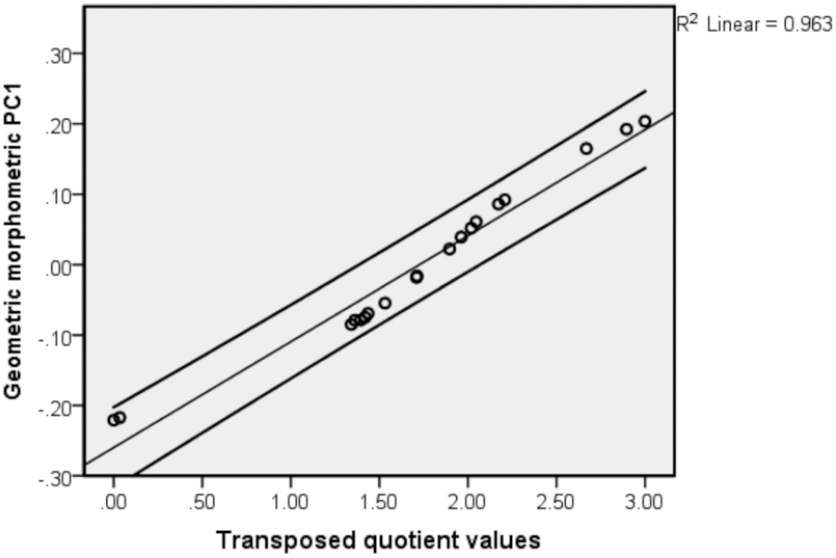
A bivariate plot of PC1 and LTS values taken from collinear data of pterosaur ulnae and wing-metacarpi. The data has a strong curvilinear correlation, but it is close to linear with a Pearson’s correlation coefficient of 0.98 and a R^2^ value of 0.96.

#### Method two

In the second experiment, cladistic analyses were run using no weighting procedures (i.e. equal weights), but with morphometric characters transformed using LTS, trigonometric and log ratio transformations. Each analysis, using each method was resampled twelve times with different combinations of dividends and divisors being used. One of the resampled replicates (matrix 1) represents the original dividend/divisor relationships of Vidovic and Martill (2014), another replicate uses the inverse (matrix 12), and the ten additional replicates are randomly resampled (*see* Supplementary Material). All 12 trees from each method were tested for their similarity using a consensus based metric, the clade retention index (CRI) (Vidovic and Martill 2017). The CRI is similar to the consensus fork index (CFI) (Colless 1980), but it accounts for the polytomous taxa, and therefore how much information is retained by a consensus tree calculated from two fundamental trees. Like many tree similarity metrics (Colless 1980; Robinson and Foulds 1981; Estabrook et al. 1985; Nixon and Carpenter 1996; Wheeler 1999; Goloboff 2008) the CRI examines a different aspect of tree similarity to others in the literature, and used alongside another method its results can be better interpreted. Here, it is used alongside tanglegrams, SPR distances (Goloboff 2008), and the R-F symmetric sampling method (Robinson and Foulds 1981) (Table 1).

#### Log ratio transformation

Mongiardino Koch et al. (2015) were not explicit regarding their methodology, but it is assumed they did not use the natural log to scale their quotient values. Therefore, here *i=log(a/b)*, where log to the base ten is used to scale the data, *i* is the index number analysed, *a* is the dividend and *b* is the divisor. During resampling *a* and *b* of the primary matrix may be inverted to give *i=log(b/a)*. The index number *i* is normalised so that all positive and negative indices lie in the range (positive) naught to one, using the range scaling equation *1/((R*_*max*_*-R*_*min*_*)x(i-R*_*min*_)), where *R* is the range.

#### Trigonometric transformation

Full details of the trigonometric scaling method can be found in Vidovic and Martill (2014). Unlike the original analysis *i=tan*^*-1*^*(a/b)* is applied to every continuous character with a quotient value, to demonstrate that it can eliminate researcher decision as well as the effect of converse datasets. Like log ratio scaled data, here the trigonometric ratio scaled data is normalised between naught and one.

#### Linear transposition scaling (LTS)

This new method implements the equation *i=(a/b)-1* if a/b>1, or *i=-((b/a)+1)* if a/b<1. In the equations, one is subtracted or added respectively, to realign the data so that it is ‘centred’ on naught. As with the former transformation methods, the LTS dataset is range scaled between naught and one using the equation given in the log ratio section of this methodology. By range rescaling the data, the weights implied by the continuous data will have a similar effect on the tree search as discrete characters (Pereyra and Mound 2009).

#### Method three

The intention of the third method is to demonstrate that quotient values maintain morphometric information when properly transformed. The results of geometric morphometric principal components analyses were compared to the results of LTS, using a bivariate plot, Pearson’s correlation coefficient and R^2^ values. The most significant principal components were chosen from a scree-plot to compare to the LTS. A geometric morphometric generalized Procrustes analysis was conducted using the Geomorph package in R (Adams et al. 2015) on real data. Bitmap images of three nodes that represent the proximal and distal ends of pterosaur ulnae and wing metacarpi (Fig. 1) were digitized in TPSdig2 (Rohlf 2010). The Bitmaps were drawn based on the exact same measurements that were used to calculate the LTS values for character 5 (see supplementary data), taken from Vidovic and Martill (2014). The ulnae and wing-metacarpi needed to be redrawn in a straight line, so that no inferences of shape can be made by their position of preservation. This practice makes the data collinear, meaning that it cannot be rotated (Small 1996). To test examples that can be rotated and rescaled in a geometric morphometric analysis, triangles (i.e. three landmarks per specimen) drawn using the dimensions of the pterosaur rostral index (Vidovic and Martill 2014: Ch.1) were analysed using the same generalized procrustes geometric morphometric method. Like with the one-dimensional material the significant PCs were identified using a scree plot and were then plotted against the LTS values in bivariate plots and in a three-dimensional scatter plot.

## Results

### Rationale/Proof of Concept

The untransformed quotient values (Fig. 3A) demonstrate both tenets of criteria 1 in continuous states calculated where the dividend is greater than the divisor, but this is not the case for continuous states calculated where the divisor is greater than the dividend. Therefore, not transforming quotient values violates criteria 2.

Both trigonometric (Fig. 3B) and logarithmic (Fig. 3C) transformations of quotient values satisfy criteria 2, but violate both tenets of criteria 1. The LTS method (Fig. 3D) of transforming quotient values satisfies both criteria 1 and 2. Thus, LTS has the most desirable effects on the transformed data.

### Method One

The graphical comparison of the rostral index (Vidovic & Martill 2014: Ch. 1) shows that when no data transformation is used and the dividend and divisor are inverted, the quotient values fail to exhibit the same morphological variation (Fig. 4A). The inequality of morphological variation observed between a character and its inverted form is because when the dividend is larger than the divisor there is a linear relationship between the source data and output, but when the divisor is larger than the dividend the relationship becomes exponential (Fig. 4A). Thus, the same variation observed between one and infinity can be demonstrated between naught and one. However, when all three transformation methods are applied, quotient values exhibit the same morphological variation for one equation (i.e. rostral depth/length) and its inverse form (i.e. rostral length/depth) (Fig. 4B-D). This practically demonstrates quotient values that are not transformed violate criteria 2.

The univariate plots demonstrate the compression and rarefaction in the logarithmic and trigonometric transformed data that is more spread or more tightly associated respectively in the untransformed data. This indicates that those transformations would not accurately represent (weight) the morphological information in a cladistic analysis.

### Method Two

Eleven of the resampled cladistic analyses for each data transformation method produced trees with SPR distances of 1.000 with respect to their reference tree (MPT from matrix 1). When comparing trees from each scaling method to trees from another, they had congruent leaves, but internal nodes shifted depending on the method used. The tree produced by the trigonometric transformation method is a strict consensus of two MPTs and is the least similar to the other two trees (Table 1). The SPR distance is limited in comparison to CRI, because it considers both logarithmic transformation comparisons to the other two methods to be equally similar, even though the other two comparison methods contradict it.

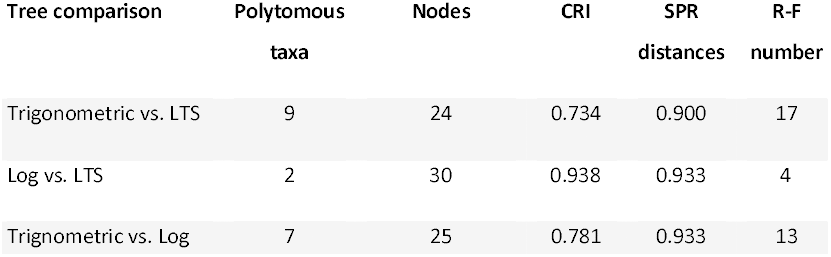

| Tree comparison | Polytomous<br>taxa | Nodes | CRI | SPR<br>distances | R-F<br>number |
| --- | --- | --- | --- | --- | --- |
| Trigonometric vs. LTS | 9 | 24 | 0.734 | 0.900 | 17 |
| Log vs. LTS | 2 | 30 | 0.938 | 0.933 | 4 |
| Trigonometric vs. Log | 7 | 25 | 0.781 | 0.933 | 13 |

### Method Three

Because the three digitized landmarks representing the terminal ends of ulnae and wing metacarpi are linear it is not possible to rotate the samples in morphospace (Small 1996: pp. 14–16). Therefore, the only principal component (PC) that was informative was PC1, which is demonstrated on the scree-plot in the supplementary data. Only having one informative PC after a generalized Procrustes analysis is not ideal for a geometric morphometric study, but in this case the data is easier to interpret and compare to the LTS method. PC1 was compared to LTS values calculated from the same specimens were used in the geometric morphometric PC analysis. The resulting bivariate plot (Fig. 5) demonstrates that the data fits a linear regression line well, with a Pearson’s correlation coefficient of 0.98 and R^2^ value of 0.96. Therefore, each method produces results that are a good approximation of one another. The reason for the output data of the two methods not being directly linear is that the geometric morphometric method projects the data into tangential morphospace, causing the curvilinear data distribution seen in the bivariate plot (Fig. 5). Note, the curve is centred on 0.00 of PC1.

The principal component analysis resulting from the generalized procrustes analysis of the pterosaur rostrum demonstrated that pterosaur rostra occupied a large continuum of morphospace (Fig. 6). PC 1 was the most significant PC, while PC 2 still demonstrated some information about morphospace. Both PC 1 and PC 2 demonstrate a well correlated curvilinear relationship with the LTS values. This means that each method can predict the other, but they share a complicated mathematical relationship as a result of projecting the shape data into morphospace during the geometric morphometric analysis.

**Figure 6.**
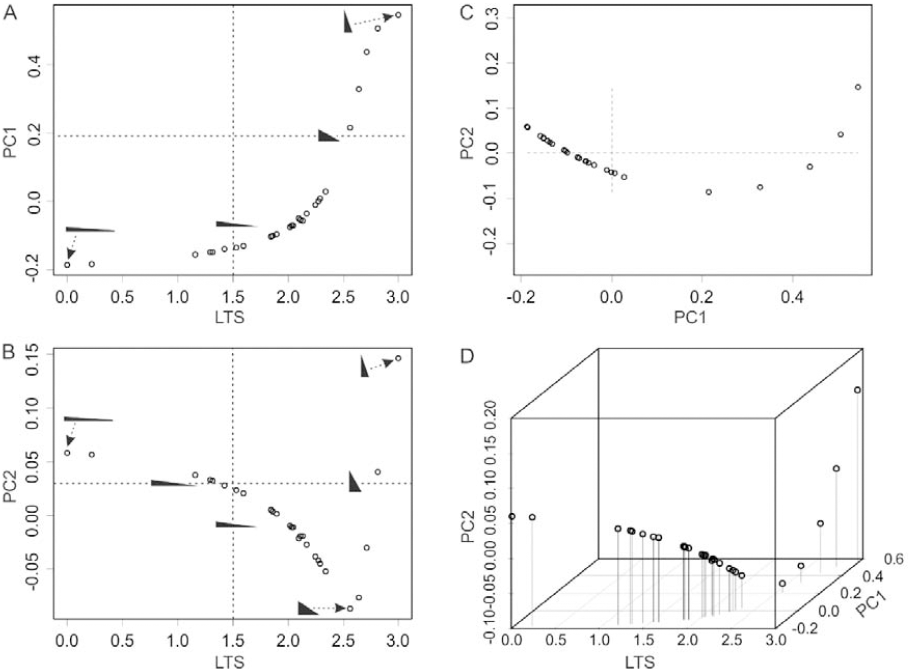
PC1 and PC2 compared to the LTS for two-dimensional shape, taken from the rostral index of Vidovic and Martill (2014). Bivariate plots of A) PC1 against LTS; B) PC2 against LTS, the triangles with arrows represent the rostrum shapes at the extremes of each range, and triangles on the dashed lines are the centre of the ranges. C) The principal component plot of PC1 and PC2; D) a comparison of LTS to both principal components in a three-dimensional plot.

Each variable on the three-dimensional plot figured is informative, especially in combination, but continuous cladistic characters can only interpret data on a univariate scale by applying weights to the states. This is also true of historic methods such as gap weighting and finite mixture coding.

In PC 1 there is a curve very similar to that seen in logarithmically transformed data (Fig. 6A). The lowest long triangle and tallest short triangle are found at the extremes of the PC1 range, while a triangle of near equal dimensions is in the middle. In PC 2 there is quadratic relationship, meaning that pterosaurs with tall short rostra will be considered related to pterosaurs with low long rostra, whilst a pterosaur rostrum exhibiting close to equal proportions is at the other extreme of the axis. Because PC2 has a quadratic relationship there are two triangles on the centre line (Fig. 6B). The LTS values separate tall, short forms and low, long forms, while placing a triangle with a reasonably low and long form in the middle of the continuum. Considering that one extreme is significantly lower and longer than the other is tall and short, the skew towards low and long is expected. Using the LTS method also means that a similar degree of variation is weighted similarly at any point in the data continuum.

## Discussion

The comparison of quotient values for systematics assumes a constant allometry (at least for sub-mature to mature individuals), a relatively tight fit of the residuals to that allometry, and the intersection of the regression line is at naught. However, the vast majority of species diagnoses, qualitative cladistic characters and the other methods of entering continuous data also make the same assumptions. If the population variation is negligible relative to the variation observed between species, genera or higher taxonomic grades, there should be little or no overlap and the character will be phylogenetically informative. Because the underlying assumptions are not testable for most taxa in the fossil record, it is recommended that this method is used alongside implied weighting (Goloboff 1993; Goloboff et al. 2008a; Goloboff 2014).

Thiele (1993) proposed that logarithms should be exploited, because 10 mm variation between plant taxa with mean leaf lengths of 5 mm and 10 mm is significant, whereas it is not for leaves with a mean length of *c*. 100 mm. While Thiele’s (1993) observation stands for untransformed size data, it is only applicable to quotients where the divisor and dividend exhibit an obvious heteroskedastic distribution in a bivariate plot of its component measurements. Thus, prior to transformation for inclusion as a continuous character, the measurements that comprise the divisor and dividend should be log transformed so that a log/log quotient value is calculated. The results of calculating log/log quotient values are not the same as those generated by log ratio transformations on the same measurements.

The result of using log ratio and trigonometric data transformations is an equivalent distribution of morphological data, regardless of converse data or researcher decisions. However, exponential data does not represent or code the true variation exhibited by the morphology (Fig. 4A & C–D). The effect of these exponential scales is to cause rarefaction of data about the centre of the maximum possible data range, and compression of data at the extremes of the range (Fig. 3A–C). Consequently, the more ‘extreme’ in the data range two taxa become, the more extreme their variation must be to consider them unrelated. Whereas, if two very similar taxa being studied have close to equal dividends and divisors (i.e. they are approaching unity) they will be more readily distinguished if logarithmic or trigonometric data transformations are used. If data transformed using these methods is entered into a continuous cladistic character, the weights given to the states will be equivalent to this data distribution, providing a bias for splitting and lumping of taxa in different parts of the data continuum.

The LTS method effectively reduces the information from two straight line distances to a single index number, ready for its input into a cladistic analysis using continuous characters. The LTS transformation of quotient data has the least negative impact on the distribution of weights/states implied compared to the other data transformations tested, according to the logic applied here. The LTS method does this by reducing the magnitude of one dimension at a function of the other (smaller) dimension’s magnitude. When entered into a cladistic analysis using continuous characters, the LTS data will treat all morphological variation similarly by implying a weight difference between two OTUs equivalent to the difference in dimensions between those OTUs. For the study of inter-taxonomic relationships, where size has been eliminated and shape is the sole consideration, equal weighting for equal variation is important.

TNT’s continuous character function is able to manage data without excessive subjectivity – assuming there is no scale inequality, as discussed in this paper –, therefore introducing potentially confounding data as a result of logarithmic transformation, trigonometric transformation, or dividing a number by a greater number (including percentages) is undesirable. Furthermore, the use of PC data taken from geometric morphometric analyses as continuous cladistic character states would have a similar effect to using improperly transformed quotient data and could even cause completely unrelated morphologies to appear more related to each other than they are to their intermediate, transitional morphologies (Fig. 6B) if the PC demonstrates a quadratic distribution.

## Conclusions

Despite many cladistic analyses using the same specimens, the same software and broadly similar characters, there are numerous examples of different authors consistently recovering contradictory tree topologies (like Researcher 1 and Researcher 2 in the example). Notable examples in vertebrate palaeontology are the variable position of Anurognathidae and *Germanodactylus* spp. (e.g. Lü et al. 2009; Wang et al. 2009; Andres et al. 2014; Vidovic and Martill 2014) in Pterosauria, and the position and composition of Tethysuchia and/or Thalattosuchia in crocodilomorph phylogeny (Jouve 2009; Sweetman et al. 2015). Some researchers opt to test new taxa or theories against multiple phylogenies (e.g. Pol and Gasparini 2009; Holliday and Gardner 2012), not necessarily favouring one model over another, but this is arguably a pointless exercise. In these cases, one or both phylogenetic hypotheses must be highly incongruent with the exact phylogeny, so the sources of error should be identified and eliminated before proposing a more robust hypothesis. Unfortunately, it is impossible to know the exact phylogeny, so ensuring that the methods used are robust and stable is the only way to be confident in a result. If the data used in an analysis is very similar, the source of the incongruence may be the result of researcher decision and the inadvertent application of errors to different parts of the matrix. For example, it can be demonstrated that atomizing compound characters reduces character incongruence (Brazeau 2011) and reorganizes some internal nodes (pers. obs.). Likewise, the improper treatment of continuously variable characters will have a similar effect on tree topology. By using tested and rationalized character construction methods, data transformation, coding and matrix analysis, it may be possible to limit researcher decision/error and find greater consensus in our phylogenetic models.

It is evident that not scaling continuous data that demonstrates converse relationships has a negative effect on the results of a cladistic analysis (Fig. 3A & Fig. 4A), making it possible to recover entirely different trees when using the same data in different ways. However, when one measured unit consistently exceeds another throughout a dataset no data transformation is necessary, contrary to Mongiardino Koch et al. (2015), but the dividend should always be the largest number. When transformation is required it is advisable to use LTS, to avoid confounding the tree search algorithms in TNT by unevenly weighting morphological variation. However, despite the obvious problems, the logarithmic transformation does not appear to have as much of an impact on the results of a cladistic analysis as trigonometric transformation (Fig. 6). Therefore, in certain circumstances logarithmic transformations may prove useful. Likewise, the trigonometric transformation was initially developed to handle data that represents a triangle (i.e. pterosaur rostral index) and it may still prove useful to study the tangent angle in cladistic analyses when studying triangular morphologies. However, applying the trigonometric transformation to all converse data (Vidovic and Martill 2014) and the logarithmic transformation to all quotient values (Mongiardino Koch et al. 2015) studied in a cladistic analysis is improper, and in this sense the methods should generally be abandoned.

## Acknowledgements

I would like to thank the Willi Hennig Society for making the cladistic software TNT freely available. I would also like to thank Nic Minter (Portsmouth) and David Martill (Portsmouth) and David Unwin (Leicester) for discussions relating to this research.

